# Predicted templates improve the prediction of antibody-antigen structures by AlphaFold 3 and Boltz-2

**DOI:** 10.64898/2026.11.20.768078

**Authors:** Omid Ziaee, Maria Lazou, Diane Joseph-McCarthy, Sandor Vajda

## Abstract

**Motivation:** In spite of the much improved protein structure prediction methods, the success rate remains low for antibody-antigen complexes. While this can be improved by computationally demanding massive sampling, results are still much poorer than for most other types of protein-protein interactions. In principle, improvement can be achieved by using templates that are close enough to the native structure of the complex, but these are generally not available for antibody-antigen complexes.

**Results:** We explore the use of templates, generated by the computationally inexpensive rigid body docking program PIPER. Applied to a leakage-free antibody-antigen benchmark set, standard AlphaFold 3 with 1000 random seeds yields acceptable or better models as the top predictions for 56.2% of complexes. Adding the PIPER generated templates in the unmasked version of AlphaFold 3 either increases the success rate to 63.4%, or reaches the same 56.2% success rate in about 200 runs. The template-based approach similarly improves antibody-antigen prediction by Boltz-2. This result demonstrates that the space explored by the ML based approach can be expanded by adding templates prepared by traditional docking, compensating for the lack of evolutionary relationship and the limited availability of training data.

**Availability and Implementation:** The ClusPro server (https://cluspro.bu.edu/login.php) containing the PIPER program is free for non-commercial applications and the program is available by request (https://www.vajdalab.org/). All other study data are included in the article and/or supporting information.

## 1 Introduction

The outstanding performance of the deep learning based program AlphaFold2 (AF2) at the CASP-14 competition in 2020 fundamentally changed the landscape of protein modeling (Jumper, et al., 2021). It is now generally recognized that AF2, AlphaFold Multimer (AFM) (Evans, et al., 2021), RoseTTAFold2 (Baek, et al., 2021), the more recent AlphaFold3 (AF3) (Abramson, et al., 2024), and several similar programs such as Boltz-1 (Wohlwend, et al., 2025), Boltz-2 (Passaro, et al., 2025), Protenix (Team, et al., 2025), Chai-1 (team, et al., 2024), and HelixFold 3 (Liu, et al., 2024) can produce models indistinguishable from experimental structures for a large fraction of proteins and protein-protein complexes (Wodak, et al., 2023). However, the success rates remain substantially lower for predicting the structures of antibodies and antibody-antigen (Ab-Ag) complexes. For example, while AFM yields acceptable or better quality models as the top prediction over 70% for general protein complexes, for antibody-antigen pairs the reported success rates were only 17% (Giulini, et al., 2024), 20% (McCoy, et al., 2024), and 24% (Yin and Pierce, 2024). Although AF3 increases the success rate to around 80% for general protein-protein interactions, it remains below 50% the antibody–antigen complexes (Xu, et al., 2025). Results for other prediction programs are substantially poorer (Fromm, et al., 2026; Xu, et al., 2025).

Predicting the structures of Ab-Ag complexes is limited by two main factors. First, both AF2 and AF3 use neural network architectures with attention-based components that take advantage of the evolutionary information extracted from multiple sequence alignments (MSAs). However, antibody-antigen interactions are primarily dependent on the highly variable complementarity determining regions (CDRs) that are determined by clonal selection, expansion and somatic hypermutation during antibody maturation (Vajda, et al., 2021), and are unimpacted by evolutionary relationships. Thus, MSAs provide little information for the construction of initial conformations of antibodies. Second, the performance of machine learning (ML) methods heavily depends on the availability of training data, which can be the determining factor for prediction success. For example, the paper reporting the development of AF2 explicitly attributes much of the accuracy gains to curated, high-quality, non-redundant structures in the Protein Data Bank (PDB) and the very large number of protein sequences that can be used for building MSAs (Jumper, et al., 2021). The paper also shows significant performance degradation on poorly represented sequence families. Antibodies and antibody-antigen complexes are clearly in this category. While the PDB lists over 250,000 protein structures (about 90,000 clusters with 90% sequence identity) and about 30,000 protein complexes with distinct architectures, the Structural Antibody Database (SAbDab) includes only around 11,000 PDB structures, among them less than 2,500 being nonredundant (Dunbar, et al., 2014). Thus, the training set for the prediction of Ab structures and Ab-Ag complexes is two orders of magnitude smaller than the one for predicting protein structures and an order of magnitude smaller than for protein complexes in general.

There is strong evidence that additional sampling can improve the docking of proteins in general and Ab-Ag pairs in particular. The success of a massive sampling strategy was first demonstrated by Wallner (Wallner, 2023), who generated large numbers of AF2 predictions by using random seeds, each selecting a subset of sequences from the MSA, and thus introducing variation in the evolutionary information. Other parameters, including the number of recycling steps and the activation or not of dropouts were also varied, but these had relatively little impact (Raouraoua, et al., 2024). Running the calculations with many random seeds and selecting the models with the highest confidence substantially improved the accuracy of predictions (Yin and Pierce, 2024). Accordingly, the paper describing the development of AF3 program reported antibody-antigen docking results using 1000 random seeds (Abramson, et al., 2024), generating 5000 models and resulting in a success rate of around 60%. While the actual number depends on the particular benchmark set, several benchmark papers report similar success rates (Mirabello, et al., 2024; Xu, et al., 2025; Yin, et al., 2026).

The explanation for improving prediction accuracy by running the programs by many seeds is that the different seeds select different MSA subsets, resulting in different initial conformations generated by the Evoformer module in AF2 and the Pairformer module in AF3 (Roney and Ovchinnikov, 2022) that are then converted into atomic coordinates by the structure module in AF2 and the diffusion module in AF3. The success of this protocol confirms that variations in the MSA yield conformational diversity. However, the usefulness of selecting MSA subsets to increase conformational diversity is reduced if the starting MSA is not very informative, which is the case for antibodies. In this paper we consider an alternative approach, expanding the space of initial conformations by adding pre-generated structural templates. One problem is that for Ab-Ag complexes relevant template structures are generally not available due to the lack of evolutionarily related antibodies. The second problem is that AF3 uses templates only for individual chains and cannot use the information on the interactions among the chains in multimeric template structures. This limitation was recently removed in the unmasked version of AF3 (Mirabello, et al., 2024), referred here as AF3-unmasked.

To motivate the template-based approach we first show that using the native complex structures as templates yields near 100% success rate. While this result clearly has no practical implications, it demonstrates that both AF3-unmasked and Boltz-2 can accept templates. Thus, the question is whether we can generate good enough templates, preferably by a computationally inexpensive algorithm. The protocol we test here is obtaining antibody and antigen monomer models by either Boltz-2 or AF3, and then generating potential templates for the complex by systematic sampling on the 6D rotational-translational space using the very fast Fourier transform based rigid body docking program PIPER (Kozakov, et al., 2006). PIPER evaluates the energies of billions conformations, the low energy docked structures are clustered, and a number of cluster centers are selected to be used as templates for AF3-unmasked and Boltz-2 runs. An additional advantage of this protocol is the ability to limit the sampling to conformations that have the CDRs in the antibody-antigen interface.

We have previously used this template-based prediction strategy in the CASP16-CAPRI experiment, where the hybrid pipeline performed very well for the CAPRI target protein multimers, with particularly good results for antibody–antigen and viral–host interactions (Ashizawa, et al., 2026). However, CASP had only a limited number of such targets, and the present paper reports the first larger scale evaluation of antibody-antigen docking with templates generated by systematic sampling. We will also investigate the relationships between the templates and the resulting conformations. As will be shown, while the gain in the success rates is moderate, computational efforts can be substantially reduced by using the templates. We first describe the approach in the context of structure prediction by AF3, and then apply it to Boltz-2. For comparison we also tested complex structure prediction by AlphaFold Multimer (AFM).

## 2 Methods

### 2.1 Antibody–antigen benchmark dataset construction

Antibody–antigen complex structures were retrieved from the Structural Antibody Database (SAbDab), using a summary file downloaded on September 11th, 2025 (Dunbar, et al., 2014). File entries were first filtered to retain only those with defined heavy chain, light chain, and antigen chain annotations, and non-protein antigens (such as nucleic acids) were excluded.

Duplicate PDB entries were removed, retaining one record per unique PDB ID. Only structures deposited after January 1, 2024 were retained, corresponding to structures deposited after the Boltz-2 validation cutoff date, yielding 1,849 complexes.

To make sure the benchmark set contains no complexes that closely resemble structures in the Boltz-2/AF3 training or validation sets, we screened each candidate for structural redundancy against a reference set of all SAbDab antibody–antigen units released before January 1, 2024 (11,725 units from 6,050 PDB entries). Each reference unit was trimmed to its heavy chain, light chain, and antigen chains. We compared benchmark complexes to the reference set with Foldseek-Multimer (https://doi.org/10.1038/s41592-025-02593-7). We chose it because it compares 3D structure rather than sequence, and it fits all chains with a single rigid superposition, so only complexes that share the same binding pose score highly. For each benchmark complex, the top 20 Foldseek-Multimer hits were re-scored with DockQ (https://doi.org/10.1371/journal.pone.0161879), with the antibody (heavy plus light chain) treated as one partner and the antigen as the other, so the score reflects the similarity of the interface itself. A benchmark complex was removed if any hit had DockQ ≥ 0.49. It was also removed if a hit had DockQ between 0.23 and 0.49 and either an interface RMSD ≤ 2 Å or a ligand RMSD ≤ 5 Å. Complexes whose hits all scored DockQ < 0.23 were kept. This step reduced the set to 121 complexes. In a second pass, the same procedure was applied all-vs-all within the benchmark set to remove internal redundancy. Within each redundant pair, the complex with the better resolution was kept. Finally, only complexes resolved at 3 Å or better were retained, resulting in 112 complexes.

### 2.2 Structure generation

#### 2.2.1 Overview of the pipeline

Our goal is to improve antibody–antigen complex prediction by combining multi-seed sampling in AF3-unmasked with templates generated by the protein–protein docking program PIPER. The templates diversify the starting points of prediction with physically plausible binding modes, expanding the conformational space that AF3 samples. The outline of our combined docking algorithm is shown in Fig. 1. Given the sequences of the interacting proteins (Fig. 1A), an MSA is generated for each (Fig. 1B), and separate models for the antibody and the antigen are obtained using either Boltz-2 or AF3 (Fig. 1C, see section 2.2.2). The highest ranked models of the two proteins are docked to each other using PIPER and the top 10,000 models of the complex are retained (Fig. 1D, section 2.2.3). These models are clustered using 3 Å pairwise RMSD as the clustering radius to form up to 1000 clusters (Fig 1E). The cluster centroids are then used as templates in AF3-unmasked, each run producing a single model per template (Fig. 1F, section 2.2.4). This protocol will be referred to as AF3+PIPER. In parallel we also run standard AF3 with 1000 random seeds to model the antibody-antigen complex as in the original AF3 paper (Abramson, et al., 2024), generating 5 models in each run (Fig. 1G, section 2.2.5). The models obtained with the PIPER generated templates in step F and the ones obtained by standard AF3 in Step G are pooled and ranked by confidence level (Fig.1H). This protocol will be referred to as AF3+PIPER with AF3. A similar pipeline is used using Boltz-2 rather than AF3-unmasked. The pipeline includes predicting the complexes using Boltz-2 alone (section 2.2.6) and using Boltz-2 with PIPER-generated models of the complex as templates (section 2.2.7). The models obtained using Boltz-2 and the PIPER-Boltz-2 combination are pooled and ranked. All pooled models are also re-scored and re-ranked using both AF3Score and ABAG-Rank (section 2.2.8) (Liu, et al., 2025; Tadiello, et al., 2026). Finally, for comparison we also generate complex models using the Colabfold implementation of AFM (section 2.2.9) (Mirdita, et al., 2022).

**Fig 1.**
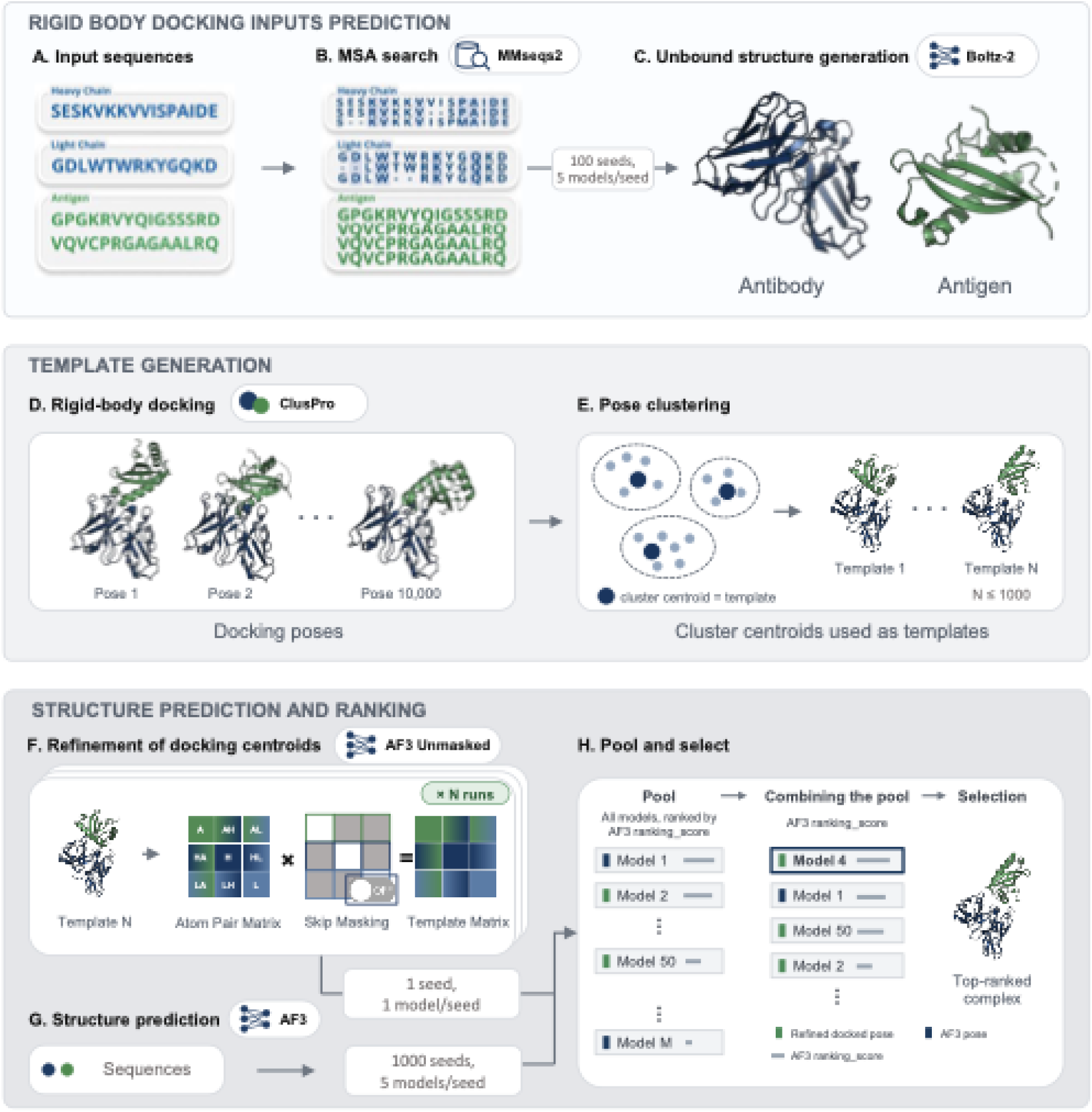
Outline of the proposed docking algorithm. (A) The sequences of the interacting proteins are used as the input. (B) An MSA is generated for each sequence. (C) Models of the component proteins are generated using Boltz-2. (D) The highest ranked models for the subunits are docked to each other using PIPER implemented in the ClusPro server, and top 10,000 poses are selected. (E) The poses are clustered and up to 1000 cluster centroids are selected. (F) Cluster centroids are used as templates for AF3-unmasked prediction, referred as AF3+PIPER. (G) In addition, standard AF3 is used to predict 1000 seeds with sequences from step A. (H) Poses from steps G (AF3+PIPER) and H (standard AF3) are pooled and ranked resulting in the combined method AF3+PIPER with AF3.

#### 2.2.2 Generating models of unbound antibody and antigen

Monomer antibody and antigen models were predicted using either Boltz-2 v2.2.1 or AF3 v3.0,0. For each benchmark case the antibody (paired heavy and light chains) and the antigen were supplied as separate inputs and predicted independently rather than as a single complex using 5 diffusion samples per seed and built-in MSA search. 100 seeds used per target, resulting in 500 models. For each component, the model with the highest ranking confidence score was retained and used as the unbound input structure for rigid-body docking.

#### 2.2.3 Prediction of complex structures to be used as templates

The unbound antibody and antigen models where used as inputs to docking by PIPER. PIPER is an FFT-based rigid-body docking program that evaluates a scoring function of shape complementarity (attractive and repulsive van der Waals terms), electrostatics (Coulomb and Born solvation models), and the knowledge-based pairwise structural potential DARS for antibody–antigen interactions (Kozakov, et al., 2006). Antibody receptors were first renumbered to the Chothia scheme, then passed through the PIPER antibody preprocessing wrapper, which produces the minimized receptor and ligand structures together with a CDR mask. Docking used the antibody parameters. Attractive van der Waals interactions were suppressed for all receptor atoms within 1.0 Å of a masked atom, biasing sampling toward the CDR paratope. The top 10,000 poses by PIPER score were clustered using greedy clustering with 3.0 Å clustering radius. Clusters were then ranked by population and represented by their model at the centroid.

#### 2.2.4 Prediction of complex structures using AF3-unmasked with PIPER generated structures as templates

PIPER cluster centroids were refined with AF3-unmasked built from source and run with the released model parameters. Each centroid was combined with the antibody receptor into a complex and supplied to as a structural template with explicit query–template residue index mapping. The multiple sequence alignments generated by the AlphaFold 3 data pipeline for each case were reused unchanged, and inference was run with the data pipeline disabled, so the docked pose was the only input that varied between poses. One model was produced per template. As mentioned, this algorithm is referred to as AF3+PIPER.

#### 2.2.5 Prediction of antibody-antigen complexes using standard AF3

Complex models were also generated using standard AF3 v3.0.0. We used 1000 seeds per complex using default model parameters (5 diffusion samples per seed), resulting in 5000 models per benchmark case. The models with the Top 1, 5 and 10 highest ranking confidence score were selected as the structures for downstream analysis.

#### 2.2.6 Prediction of complexes using Boltz-2

Models of the complexes were generated using Boltz-2 v2.2.1 (Passaro, et al., 2025) with 1000 seeds per complex, using 5 diffusion samples per seed and built-in MSA search, resulting in 5000 models per benchmark case. The models with the Top 1, 5 and 10 highest ranking confidence score were selected as the structures for downstream analysis.

#### 2.2.7 Complex structure prediction with Boltz-2 using PIPER generated templates

Templates were passed under Boltz-2’s default template handling and were not forced during inference, so the model was informed by, but not constrained to follow, the supplied structures. All other settings, including 1000 seeds per complex, 5 diffusion samples per seed and built-in MSA search, were unchanged, resulting in 5000 models per benchmark case. The models with the Top 1, 5 and 10 highest ranking confidence score were selected as the structures for downstream analysis.

#### 2.2.8 Rescoring Boltz-2 Predictions

For each benchmark complex, all models generated with Boltz-2 were rescored utilizing either the AF3score (Liu, et al., 2025) or the The ABAG-Rank (Tadiello, et al., 2026) algorithms.

#### 2.2.9 Complex structure prediction using Colabfold

ColabFold v1.6.1 was used to predict complex structures. MSAs were generated by the ColabFold MMseqs2 server. Predictions were obtained with 1,000 seeds per complex and five models per seed (5,000 models per benchmark case), using three recycles, no structural templates and no Amber relaxation. The models with the Top 1, 5 and 10 highest ranking confidence scores (0.8 × ipTM + 0.2 × pTM) were selected as the structures for downstream analysis.

### 2.3. Assessing prediction results with DockQ

Docked models were scored with DockQ (Basu and Wallner, 2016) using the original reference implementation (DockQ.py v1 with its compiled fnat helper). Since PIPER runs provide only the variable positions of the antigen whereas the antibody structure is fixed, each cluster centroid was first merged with the antibody structure in PyMOL to form a complex model; AF3 and Boltz-2 models were assessed as predicted. The reference structure was assembled in PyMOL from the X-ray antibody and antigen chains of each complex in the benchmark and standardized before comparison, sequential residue renumbering, removal of insertion codes, and retention of a single alternate location, after which its chain identifiers and residue numbering were mapped onto those of the model (via DockQ’s fix_numbering.pl global alignment) so that native and model chain groups correspond. DockQ was then run per model as DockQ.py <model> <native> -model_chain1 "H L" -perm2, i.e. the antibody heavy and light chains were grouped as interaction partner 1 and the remaining (antigen) chains as partner 2, with all antigen-chain permutations tested and the mapping giving the highest DockQ retained. Default settings were used otherwise, ranging from 0 to 1. Fnat, Fnonnat, iRMS, LRMS and DockQ were recorded for every model (16-way parallel), and models were assigned CAPRI quality classes using the standard DockQ thresholds of 0.23 (acceptable), 0.49 (medium) and 0.80 (high).

## 3 Results

### 3.1 Complex structure prediction with native structure as the template

We first have tested whether AF3, AF3-unmasked, and Boltz-2 can accept native complex structures as templates following the protocols described in the Methods, except that the bound crystal complex of each case was supplied as a single structural template (for Boltz-2 under its default template handling, i.e. not forced during inference), and used 10 seeds, giving 50 models per case. A prediction was considered successful if its DockQ score was ≥ 0.23, corresponding to CAPRI’s acceptable quality or better; success rates are reported for the top-1, top-5 and top-10 ranked poses, and for the oracle (all poses pool). As shown in Fig. 2A, the acceptable or better success rates for both Boltz-2 and AF3-unmasked are excellent, the latter reaches 99.1% for the top prediction. Fig. 2A also demonstrates that standard AF3 can take only limited advantage of the templates, although the 72.3% rate indicates that the templates still have some impact. As shown in left panel of Fig. 2A, with the native templates Boltz-2 and particularly AF3-unmasked both mostly yield medium or high accuracy top models and for AF3-unmasked a very large fraction (80.4%) of the top models are of high accuracy. Note that using the native template AF3-unmasked not only produces high quality models, but these are also ranked as Top-1 predictions. While these results have no practical significance, they clearly demonstrate that both AF3-unmasked and Boltz-2 provide excellent predictions if high quality structures are available as templates.

**Fig 2.**
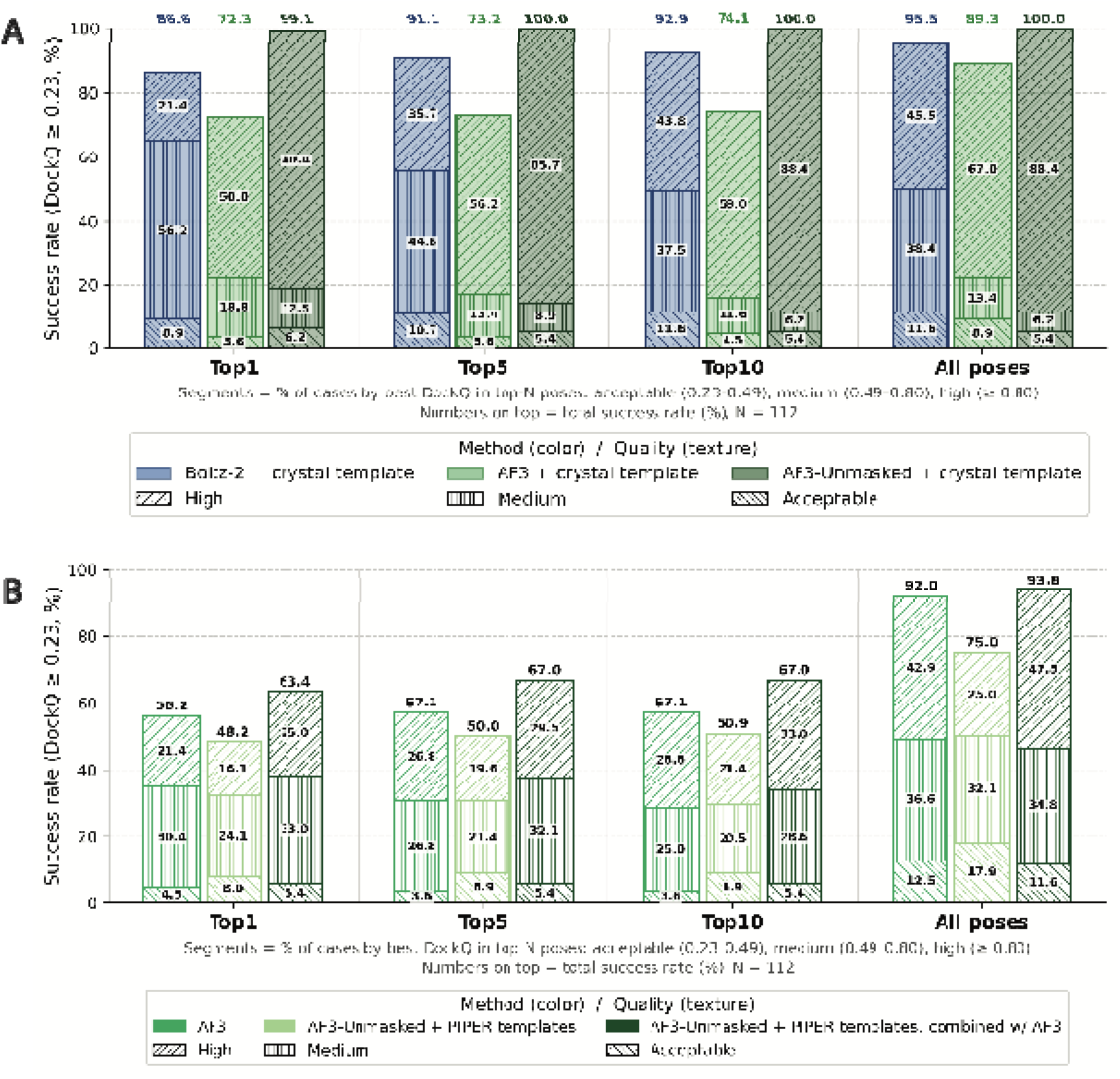
Success rates on the benchmark of 112 antibody–antigen complexes with complex structures as templates. (A) Success rates of Boltz-2, standard AF3, and AF3-unmasked with native X-ray complex structures as templates. Success is defined as DockQ ≥ 0.23. Rates are shown for the Top-1, Top-5 and Top-10 ranked poses and for all poses, stacked by CAPRI quality class defined as acceptable (0.23 ≤ DockQ < 0.49), medium (0.49 ≤ DockQ < 0.80) and high (DockQ ≥ 0.80). (B) Success rates of standard AF3, AF3+PIPER, and AF3+PIPER with AF3.

### 3.2 Combining AF3+PIPER with standard AF3 outperforms standard AF3 alone

As a baseline, we ran standard AF3 with 1,000 seeds, following the protocol used for antibody-antigen benchmarking in the paper reporting AF3 (Abramson, et al., 2024). Fig. 2B shows the success rates obtained using the standard AF3, AF3-unmasked with PIPER generated templates (AF3+PIPER), and the by pooling the results from the two previous methods, thus AF3+PIPER with AF3. AF3-unmasked with PIPER templates (AF3+PIPER) alone underperformed standard AF3 (top-1 success 48.2% vs 56.2%), yet pooling the two sets raised top-1 success to 63.4%, indicating that docking templates recover binding modes that AF3 misses. The improvement was driven mainly by higher-quality predictions: the fraction of targets with medium and high-quality top-1 poses increased from 51.8% for standard AF3 to 58.0% for the combination, whereas acceptable-quality predictions increased by only 0.9% (Fig. 2B). Fig. 2B also shows the success rates when considering all poses, which are very high (92.0% and 93.8%, respectively) for standard AF3 and AF3+PIPER with AF3. Comparing this to the top-1 success rates reveals that while good models are generated by both protocols, ranking is still a problem, and that adding the templates results in additional high-quality models that are also ranked as the best ones.

### 3.3. Combining AF3+PIPER with standard AF3 reaches AF3’s accuracy with the fraction of the sampling

Because pooling baseline AF3 and AF3+PIPER predictions increases the total number of models, we asked whether the improvement simply reflects more sampling. We therefore compared top-1 success as a function of the number of models generated (Fig. 3A) and the number of runs (Fig. 3B), adding seeds or templates. For this comparison we assumed that each AF3 run (five models that we will call here samples) is followed by two AF3+PIPER runs, each resulting in one additional model, thus each three AF3 run adds seven new samples. As shown in Fig. 3A, in this experiment the success rate of combining AF3+PIPER with AF3 reaches the maximum success rate of standard AF3 (56.2%) in only 30 runs. However, selecting different sets of runs we can also calculate upper and lower thresholds on the success rates. Considering the low threshold, the AF3+PIPER with AF3approach matched the maximum top-1 success rate of AF3 after only 202 runs (68 AF3 and 134 AF3+PIPER runs), resulting in 470 models, whereas AF3 alone requires up to 1000 runs to reach this level of accuracy when accounting for the low success rate threshold (Fig. 3A). Thus, the proposed combination of methods results in a five-fold reduction in the number of total AF3 runs. With further sampling, the combined approach continued to improve the success rate, plateauing at 63.4%. Thus, the gain from docking templates reflects more informative sampling, allowing for reduced computational efforts to reach the maximum success rate of standard AF3.. Top-1 success for every combination of the numbers of AF3 and AF3+PIPER runs, and for different ratios of AF3+PIPER to AF3 runs, is given in Supplementary Table S1 and Supplementary Fig. S1.

**Fig. 3.**
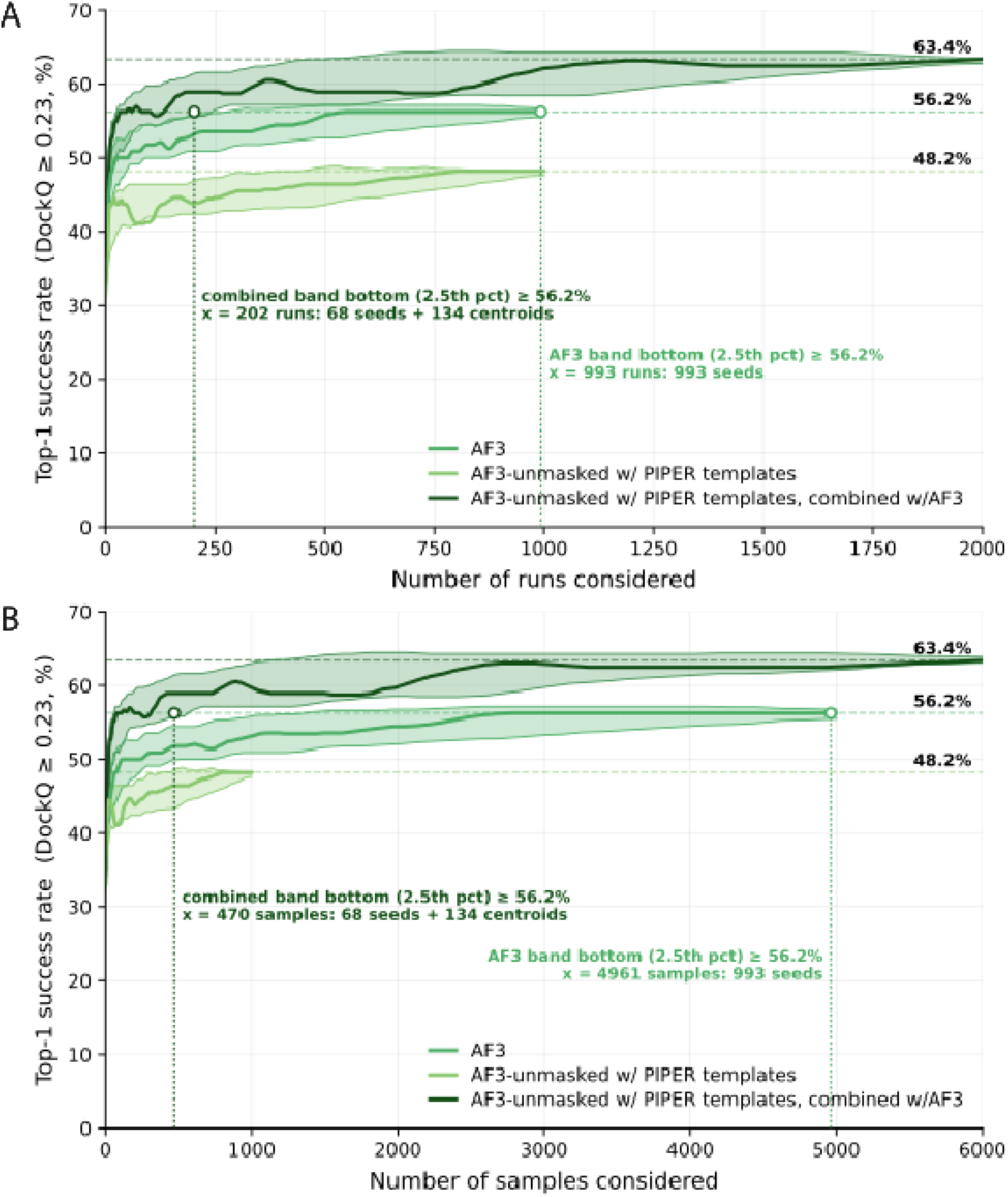
PIPER templates reduce required efforts. (A) Top-1 success rates as functions of the number of runs for standard AF3, AF3+PIPER, and the combined AF3+PIPER with AF3. In the combined method one AF3 run is combined with two AF3+PIPER runs. At each step the Top-1 model is selected by AF3 ranking score. Dashed lines mark the final Top-1 success rates of each method. Shaded regions indicate 95% confidence intervals from 100 bootstrap resamples. Dotted lines mark the point at which the lower threshold for each method first matches the maximum success of baseline AF3, 56.2%. (B) Top-1 success rates as functions of the number of samples (models). Each AF3 run yields five samples and each AF3+PIPER run yields one sample. Thus, with the selected ratio of one AF3 and two AF3+PIPER runs, in the combined method AF3+PIPER with AF3, three AF3 runs yield a total of seven samples (five from one AF3 and two from the two AF3+PIPER runs).

### 3.4 PIPER templates rescue AlphaFold3 failures

To see where the gains come from, we compared the DockQ values of the top-1 predictions by the combined AF3+PIPER and the standard AF3 target by target (Fig. 4A, left). We call a complex rescued if AF3 fails (Top-1 DockQ < 0.23) but AF3+PIPER succeeds (DockQ ≥ 0.23) and lost when reversed. On its own, AF3+PIPER rescued 11 complexes but lost 20, and its DockQ scores correlated only moderately with those of AF3 (Spearman ρ = 0.56). The two methods therefore fail on largely different targets. Generally, the targets each method fails on are of different interface difficulty class, however, PIPER-AF3 achieves higher success rates on it’s own for rigid interface complexes (Supplementary Fig. S2). Pooling AF3+PIPER with AF3 kept all 11 rescues and cut the losses to 3 (ρ = 0.85; Fig. 4A, right). The combined method gave a higher DockQ than AF3 for 43 complexes and a lower one for 34. The rescued complexes spanned all interface difficulty categories, including difficult ones. For each of the 11 rescued complexes, we compared the DockQ of the PIPER template (cluster centroid) with that of the final Top-1 AF3+PIPER model (Fig. 4B). The final model had a higher DockQ than its template in 10 of 11 cases, and the two DockQ values were not correlated (ρ = -0.01). Five templates with near-zero DockQ led to high-quality models (DockQ > 0.80).

**Fig. 4.**
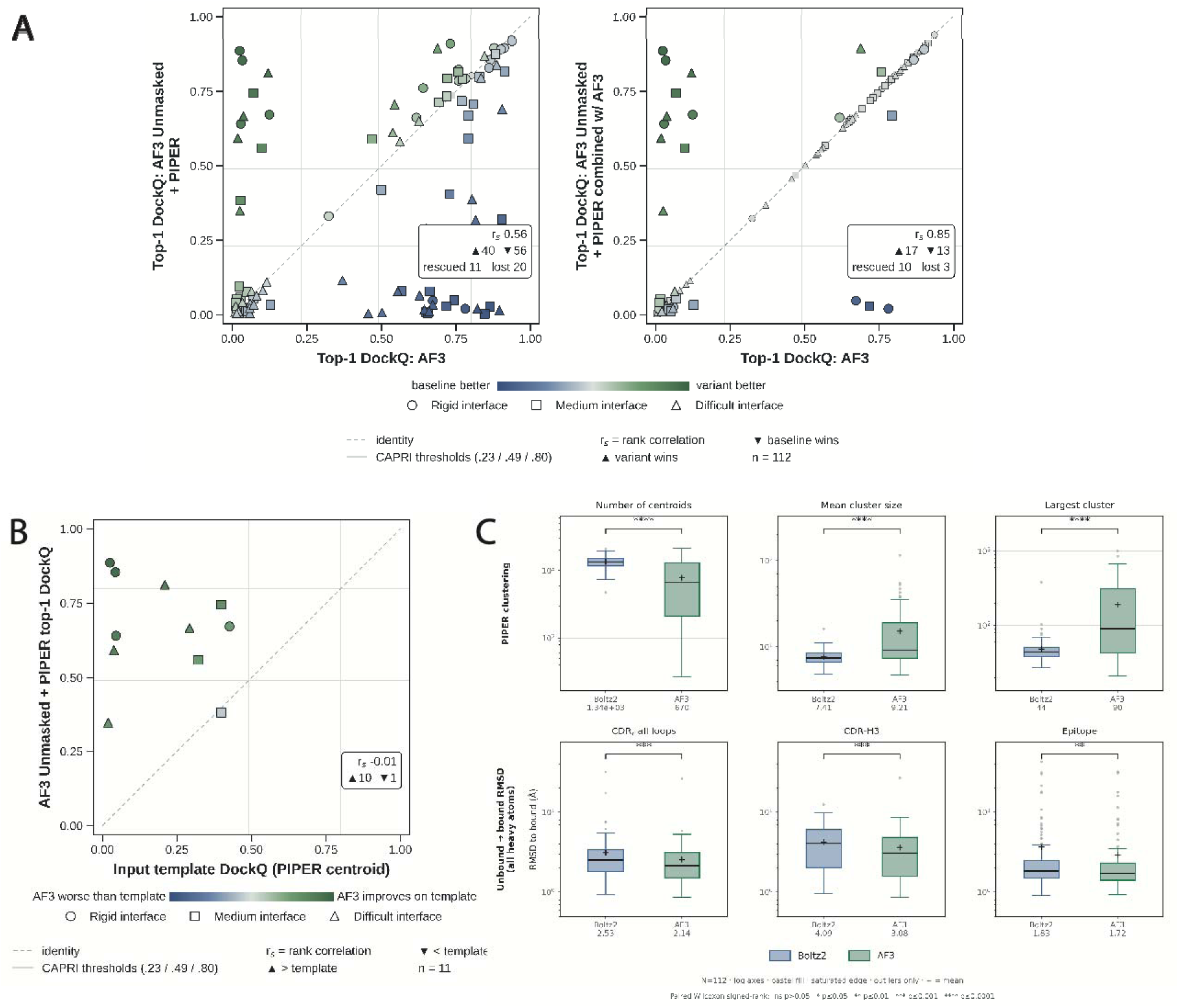
DockQ scores in the different steps of the algorithm. (A) Comparisons of Top-1 models obtained by AF3 versus by AF3+PIPER, and Top-1 models by AF3 versus AF3+PIPER with AF3. (B) DockQ scores of the 10 complexes where AF3+PIPER yields better results than AF3 versus the DockQ of the PIPER template. This shows remarkable improvement upon refinement of the input templates. (C) Impact of the models of unbound antibody and antigen models used for generating the complex templates by PIPER. Boltz2 generated unbound structures are more diverse and less similar to the bound structure, making Boltz2 structures more suitable for the starting point of the AF3+PIPER algorithm.

Because PIPER docks separately predicted antibody and antigen models, the conformations of these unbound models shape the docking results. We therefore compared PIPER docking runs starting from unbound models generated by either Boltz-2 or AF3 (Fig. 4C). Boltz-2 models deviated more from the experimental bound conformation at both the epitope and CDR loops; (2.53 Å vs 2.14 Å for CDR loops; and 1.83 Å vs 1.72 Å for epitope; both p values < 0.01; Fig. 3C, bottom). PIPER docking with Boltz-2 unbound partners also produced significantly more cluster centroids, with smaller mean and largest cluster sizes (p < 0.0001; Fig. 4C, top), which results in a broader and more diverse set of docking poses. AF3 models, being closer to the bound state, gave fewer and denser clusters dominated by one large cluster. The choice of monomer predictor is thus a trade-off between pose diversity and convergence toward bound-like poses. We use Boltz-2 models in the pipeline (Fig. 1C) because their more diverse templates give AF3 more distinct starting points, and, as Fig. 4B shows, AF3 does not require an accurate template to produce an accurate model. The pipeline was repeated with AF3-generated unbound partners for docking, which led to lower success rates (Supplementary Fig. S3 and Supplementary Table S2).

Three examples show how the combination AF3+PIPER succeeds where both AF3 and PIPER alone fails (Fig. 5). For complexes from the Protein Data Bank (PDB) - 8acf, 8zu9, and 8wsp - AF3 placed the antibody on the wrong epitope (DockQ = 0.04, 0.09, and 0.02, respectively). PIPER found roughly correct docked structures in two cases (DockQ = 0.29, 0.32, and 0.04), and with these poses as templates, AF3+PIPER produced much better models (DockQ = 0.67, 0.56, and 0.59).

**Fig. 5.**
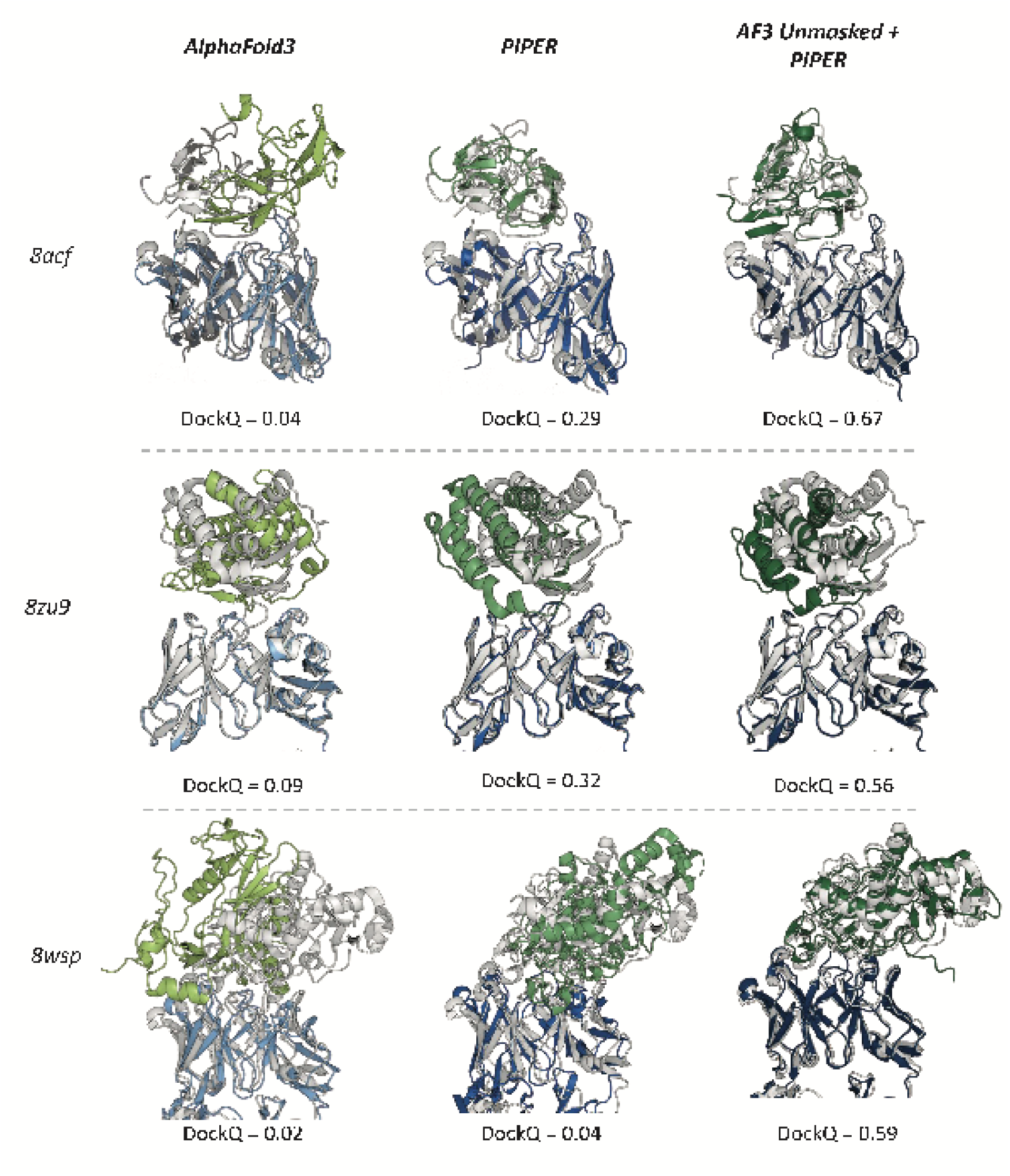
Top-1 predictions for three complexes in the benchmark. For each target we show the prediction using the standard AF3, the PIPER generated structure that was used as the template, and the result of refining the template by AF3+PIPER.

### 3.5 Templates also improve docking results by Boltz-2 when paired with effective ranking

To test whether the benefit of docking templates depends on AF3 specifically, we applied the same template-based approach to Boltz-2. Unlike AF3, Boltz-2 natively accepts multi-chain templates, allowing PIPER docking poses to be supplied directly without modifying the model. Because the complexes in the benchmark were added to the PDB after the Boltz-2 validation cutoff, we evaluated Boltz-2 on the same 112 antibody-antigen complexes. We compared four protocols: standard Boltz-2; Boltz-2 with PIPER templates; and Boltz-2 with PIPER templates re-ranked by one of two scoring methods. The first, AF3Score, applies the AF3 confidence module directly to Boltz-2-generated structures, bypassing AF3 structure generation. The second, ABAG-rank, is a DeepSets-based network trained to predict the interface quality (DockQ) of AF3-generated antibody-antigen models. As an additional reference, we included AF2.3-Multimer run through ColabFold, the most recent AlphaFold version available for commercial use.

Adding PIPER templates alone changed the top-1 success rates from 12.5% to 17.9% (Fig. 6A). Across the full pool of generated structures, however, the fraction of targets with at least one acceptable pose increased from 62.5% to 76.8%, meaning that templates enabled Boltz-2 to generate a correct pose for 17 additional targets. This gap between top-N and oracle success suggests that Boltz-2 frequently generates correct poses but fails to rank them highly. The composition of the Boltz-2 and AF3 confidence scores and their relation to DockQ are described in Supplementary Tables S3–S6 and Supplementary Figs. S4 and S6). Because all three template-based methods draw on the same pool, differences in their top-1 success arise solely from ranking. Re-ranking with AF3 score increased the top-1 success rate to 35.7%, outperforming AF2.3-Multimer (30.4%), and reached 44.6% success rate in the top-10 (Fig. 6A). Results for Boltz-2 with the single-chain templates retrieved by the AF3 data pipeline are given in Supplementary Table S7 and Supplementary Fig. S5.

**Fig 6.**
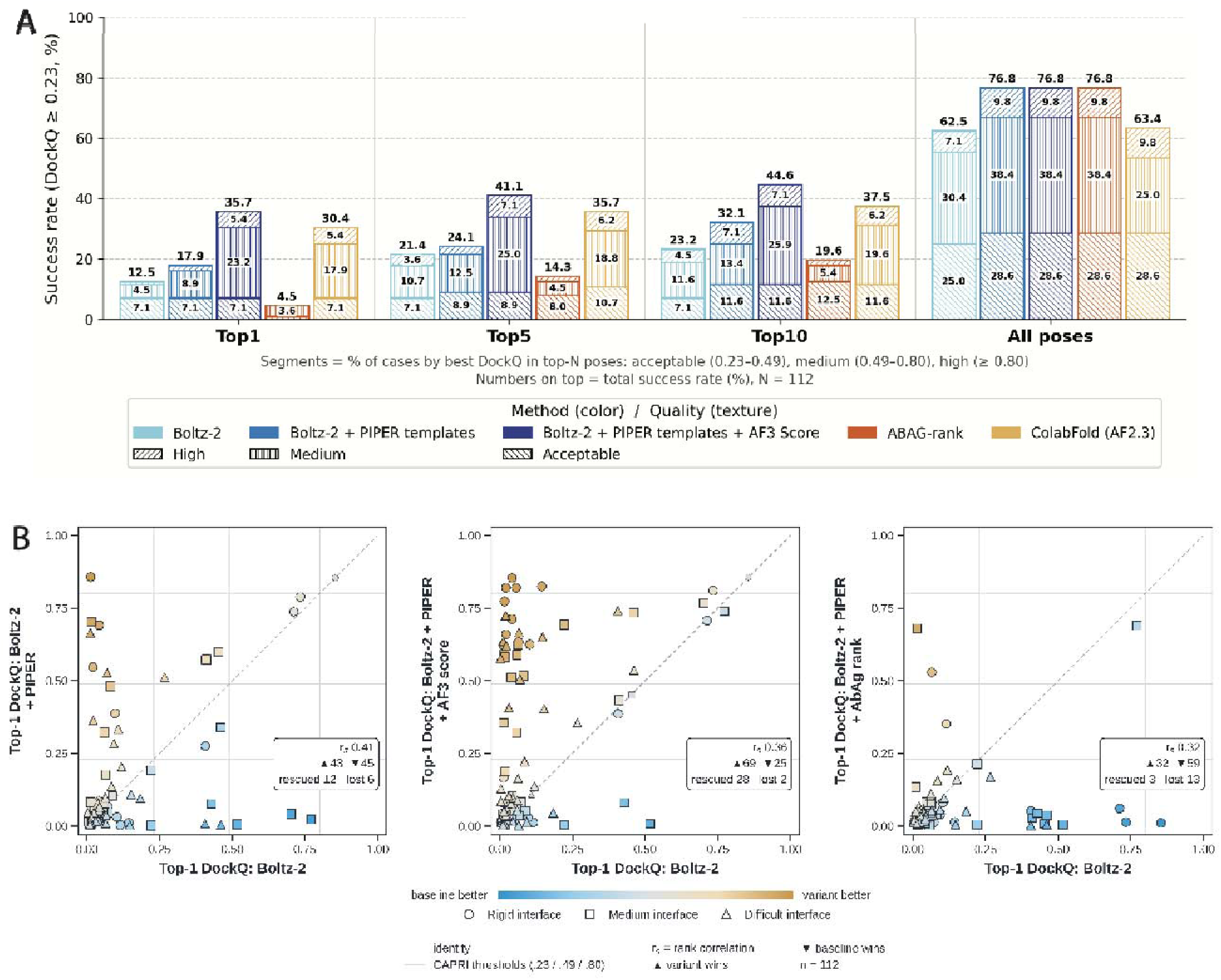
Docking templates improve Boltz-2 prediction of antibody-antigen complexes when combined with effective ranking. Results on the benchmark of 112 antibody–antigen complexes. Success is defined as DockQ ≥ 0.23 (CAPRI acceptable or better). (A) Success rates for baseline Boltz-2; Boltz-2 with PIPER templates; Boltz-2 with PIPER templates re-ranked by ABAG-rank; Boltz-2 with PIPER templates re-ranked by AF3Score; and AF2.3-Multimer, shown for the Top-1, Top-5 and Top-10 ranked poses and for oracle. Bars are stacked by CAPRI quality class: high (DockQ ≥ 0.80), medium (0.49 ≤ DockQ < 0.80) and acceptable (0.23 ≤ DockQ < 0.49). (B) Per-target Top-1 DockQ of each template-based variant against baseline Boltz-2. Points above the diagonal indicate targets improved by the variant. Marker shape denotes interface difficulty [rigid / medium / difficult (see Methods)], and color indicates the difference in DockQ between variant and baseline. ▴ and ▾ give the number of targets on which the variant or the baseline scored higher; “rescued” and “lost” give the number of targets whose Top-1 DockQ crossed the 0.23 threshold upward or downward, respectively. R_s_, Spearman rank correlation.

In contrast, ABAG-rank reduced top-1 success to 4.5%, well below standard Boltz-2, likely because it was trained on AF3-generated models and does not generalize to Boltz-2 outputs. Per-target comparisons showed the same pattern. Relative to the standard Boltz-2, templates with Boltz-2’s own ranking rescued 12 targets but lost 6 (Fig. 6B, left). Re-ranking with AF3 score rescued 28 and lost only 2 (Fig. 6B, middle), whereas AbAg-rank rescued 3 and lost 13 (Fig. 6B, right). The benefit of docking templates therefore depends critically on a scoring function able to identify correct antibody-antigen poses. For the 14 targets rescued by PIPER templates, the final Boltz-2 prediction improved on the input template in 11 cases, including templates with DockQ < 0.23 that were refined to medium-quality models (DockQ > 0.5). Final model quality correlated only moderately with template quality (r = 0.42), indicating that, as with AF3 (Fig. 4B), the network uses PIPER templates as starting points rather than copying them (Supplementary Fig. S7).

Although absolute top-1 success with Boltz-2 remained below that of AF3 (35.7% vs 63.4%), reflecting its lower baseline on this benchmark, the relative improvement by adding PIPER-generated templates was larger. Together with the AF3 results, these data suggest a general pattern: docking templates enrich the prediction pool with correct binding modes, and gains in top-1 accuracy depend on the ability to rank them. This makes the approach transferable to other template-accepting structure prediction models.

## 4 Discussion

Accurate prediction of antibody–antigen complexes remains a major challenge for deep learning structure prediction methods, largely because the MSA-derived coevolutionary signal on which these models rely is weak or absent across antibody–antigen interfaces. Increasing the number of sampled seeds partially compensates for this, but at substantial computational cost. Here, we showed that physics-based docking poses from PIPER, supplied as templates, provide an alternative route to correct binding modes. Combined with conventional seed sampling and appropriate ranking step, docking templates improved prediction accuracy over AF3, required far fewer models to match its performance, and transferred to Boltz-2 without any modifications of the model.

Our analyses point to a common mechanism behind these results. Docking explores binding modes independently of evolutionary information, and so samples regions of interface space that seed variation alone may not reach. Supplied as templates, these poses act as starting points rather than fixed answers: the networks frequently refined imperfect templates into higher-quality models, and final accuracy depended only moderately on template quality. This explains why template-guided predictions on their own were not better than baseline AF3, yet were strongly complementary to it. The two sources of sampling recover different targets, so combining them enriches the pool with more correct poses. The same enrichment underlies the efficiency gains, since a few template-guided models contribute correct poses that would otherwise require hundreds of additional seeds. It also explains the Boltz-2 results, where templates substantially increased the fraction of targets with a correct pose but improved Top-1 accuracy only once an effective scoring function was used. Across both models, then, accuracy is determined by two separable factors: whether a correct pose is generated, and whether it is ranked first. Docking templates address the first; the choice of scoring function determines the second.

Several strategies have been proposed to improve multimer prediction beyond default AlphaFold sampling. Massive sampling approaches, such as running AF3 with 1,000 seeds or AFsample (Wallner, 2023) and MassiveFold (Raouraoua, et al., 2024), increase the diversity of generated structures, but they do so by brute force and remain bounded by the model’s learned prior. Hybrid approaches have combined deep learning with physics-based docking, for example by refining AlphaFold models with ReplicaDock (AlphaRED) (Chu, et al., 2023), or by guiding prediction with experimental or predicted restraints, as in ColabDock (Feng, et al., 2024), AlphaLink (Stahl, et al., 2023) and the restraint features of Chai-1 (team, et al., 2024).

Our approach differs in passing complete docked complexes as templates, so the network receives a full candidate interface rather than sparse distance constraints, while remaining free to deviate from it. Unlike antibody-specific models, it requires no retraining and can be applied to any structure prediction model that accepts multi-chain templates. Our results also echo previous observations that confidence metrics such as ipTM are unreliable for antibody–antigen complexes (Supplementary Table S6, Supplementary Fig. S4). The poor transfer of ABAG-rank to Boltz-2 outputs further suggests that learned rankers may be specific to the model whose structures they were trained on.

Our study has several limitations. First, the benefit of docking templates depends on the docking pool containing a near-native pose. When it does not, templates can either point the network in the correct direction or do the opposite, as reflected in the targets lost relative to baseline (Fig. 4A). Improving docking sampling, for example by incorporating epitope interaction constraints may reduce these failures. Second, the largest gains required re-ranking, and the gap between Top-1 and oracle success shows that ranking remains the main bottleneck. Developing scoring functions that generalize across models, rather than being trained on the outputs of a single one, is a natural next step (Feldman and Skolnick, 2025). Third, our best Boltz-2 results relied on the AF3 confidence module for re-ranking, which is subject to the AF3 license terms. A fully permissive pipeline will therefore require an open scoring function of comparable quality. Fourth, our benchmark of 112 complexes, although constructed to avoid data leakage, does consist of only conventional antibodies and scFVs. Subsequently, performance on specific subclasses, such as nanobodies or highly flexible interfaces, warrants further evaluation. Finally, we tested the approach on AF3 and Boltz-2 only. Because it requires only template support, it should extend to other models such as Chai-1, Protenix or OpenFold3, and to other interface classes in which coevolutionary signal is limited, including T-cell receptor-peptide-MHC and host-pathogen complexes.

## Funding

This investigation was supported by the National Institute of General Medical Sciences [R35GM118078, 1R43GM161151-01]

## Notes

### Competing Interest Statement

The authors have declared no competing interest.

## References

Abramson J, Adler J, Dunger J et al. Accurate structure prediction of biomolecular interactions with AlphaFold 3. Nature 2024;630(8016):493–500.

Ashizawa R, Kotelnikov S, Khan O et al. Modeling protein–protein and protein–ligand interactions by the ClusPro team in CASP16. Proteins: Structure, Function, and Bioinformatics 2026;94(1):183–191.

Baek M, DiMaio F, Anishchenko I et al. Accurate prediction of protein structures and interactions using a three-track neural network. Science 2021;373(6557):871–876.

Basu S and Wallner B. DockQ: A Quality Measure for Protein-Protein Docking Models. PLoS One 2016;11(8):e0161879.

Chu LS, Ruffolo JA, Harmalkar A et al. Flexible Protein-Protein Docking with a Multi-Track Iterative Transformer. bioRxiv 2023.

Dunbar J, Krawczyk K, Leem J et al. SAbDab: the structural antibody database. Nucleic Acids Res 2014;42(Database issue):D1140–1146.

Evans R, O’Neill M, Pritzel A et al. Protein complex prediction with AlphaFold-Multimer. BioRxiv 2021:2021.2010. 2004.463034.

Feldman J and Skolnick J. AF3Complex Yields Improved Structural Predictions of Protein Complexes. bioRxiv 2025.

Feng S, Chen Z, Zhang C et al. Integrated structure prediction of protein–protein docking with experimental restraints using ColabDock. Nature Machine Intelligence 2024;6(8):924–935.

Fromm S, Ludaic M and Elofsson A. Evaluating deep learning based structure prediction methods on antibody–antigen complexes. Bioinformatics 2026;42(4):btag136.

Giulini M, Schneider C, Cutting D et al. Towards the accurate modelling of antibody-antigen complexes from sequence using machine learning and information-driven docking. Bioinformatics 2024.

Jumper J, Evans R, Pritzel A et al. Highly accurate protein structure prediction with AlphaFold. Nature 2021;596(7873):583–589.

Jumper J, Evans R, Pritzel A et al. Applying and improving AlphaFold at CASP14. Proteins 2021;89(12):1711–1721.

Kozakov D, Brenke R, Comeau SR et al. PIPER: an FFT-based protein docking program with pairwise potentials. Proteins 2006;65(2):392–406.

Liu L, Zhang S, Xue Y et al. Technical report of HelixFold3 for biomolecular structure prediction. arXiv preprint arXiv:2408.16975 2024.

Liu Y, Yu Q, Wang D et al. Af3score: A score-only adaptation of alphafold3 for biomolecular structure evaluation. Journal of Chemical Information and Modeling 2025;65(15):8207–8214.

McCoy KM, Ackerman ME and Grigoryan G. A comparison of antibody-antigen complex sequence-to-structure prediction methods and their systematic biases. Protein Sci 2024;33(9):e5127.

Mirabello C, Wallner B, Nystedt B et al. Unmasking AlphaFold to integrate experiments and predictions in multimeric complexes. Nature communications 2024;15(1):8724.

Mirdita M, Schutze K, Moriwaki Y et al. ColabFold: making protein folding accessible to all. Nat Methods 2022;19(6):679–682.

Passaro S, Corso G, Wohlwend J et al. Boltz-2: Towards accurate and efficient binding affinity prediction. BioRxiv 2025.

Raouraoua N, Lensink M and Brysbaert G. Massive sampling strategy for antibody-antigen targets in CAPRI Round 55 with MassiveFold. Authorea Preprints 2024.

Raouraoua N, Mirabello C, Very T et al. MassiveFold: unveiling AlphaFold’s hidden potential with optimized and parallelized massive sampling. Nat Comput Sci 2024;4(11):824–828.

Roney JP and Ovchinnikov S. State-of-the-Art Estimation of Protein Model Accuracy using AlphaFold. bioRxiv 2022.

Stahl K, Graziadei A, Dau T et al. Protein structure prediction with in-cell photo-crosslinking mass spectrometry and deep learning. Nature biotechnology 2023;41(12):1810–1819.

Tadiello M, Ludaic M, Viliuga V et al. ABAG-Rank: Improving Model Selection of AlphaFold Antibody–Antigen Complexes by Learning to Rank. bioRxiv 2026:2026.2003. 2017.712376.

Team BAAS, Chen X, Zhang Y et al. Protenix-advancing structure prediction through a comprehensive AlphaFold3 reproduction. BioRxiv 2025:2025.2001. 2008.631967.

team CD, Boitreaud J, Dent J et al. Chai-1: Decoding the molecular interactions of life. BioRxiv 2024:2024.2010. 2010.615955.

Vajda S, Porter KA and Kozakov D. Progress toward improved understanding of antibody maturation. Curr Opin Struct Biol 2021;67:226–231.

Wallner B. AFsample: improving multimer prediction with AlphaFold using massive sampling. Bioinformatics 2023;39(9).

Wodak SJ, Vajda S, Lensink MF et al. Critical Assessment of Methods for Predicting the 3D Structure of Proteins and Protein Complexes. Annu Rev Biophys 2023;52:183–206.

Wohlwend J, Corso G, Passaro S et al. Boltz-1 democratizing biomolecular interaction modeling. BioRxiv 2025:2024.2011. 2019.624167.

Xu S, Feng Q, Qiao L et al. Benchmarking all-atom biomolecular structure prediction with FoldBench. Nature Communications 2025;17(1):442.

Yin R and Pierce BG. Evaluation of AlphaFold antibody-antigen modeling with implications for improving predictive accuracy. Protein Sci 2024;33(1):e4865.

Yin R, Saravanakumar S, Shi SY et al. Benchmarking AlphaFold and related deep learning approaches for modeling antibody and TCR antigen recognition. bioRxiv 2026:2026.2007. 2004.736425.

