## Supplemental Tables and Figures for "Predicted templates improve the prediction of antibody-antigen structures by AlphaFold 3 and Boltz-2"


**Table S1.** Sampling budget at which the mean Top-1 success over 100 random draws reaches 56.2% and 62.5% when AF3 seeds and PIPER centroids are added at a fixed ratio (112 targets; models ranked by AF3 ranking score; at most 1,000 centroids). Each entry gives seeds / centroids at the first budget at which the mean reaches the target (models = 5 × seeds + centroids).

| Seeds : centroids | 56.2%: seeds / centroids | 56.2%: models | 62.5%: seeds / centroids | 62.5%: models |
|---|---|---|---|---|
| 1 : 0.5 | 43 / 22 | 237 | 711 / 356 | 3,911 |
| 1 : 1 | 31 / 31 | 186 | 603 / 603 | 3,618 |
| 1 : 1.5 | 26 / 39 | 169 | 520 / 780 | 3,380 |
| 1 : 1.9 (optimum) | 24 / 46 | 166 | 473 / 899 | 3,264 |
| 1 : 2 (Fig. 3) | 24 / 48 | 168 | 474 / 948 | 3,318 |
| 1 : 3 | 20 / 60 | 160 | 471 / 1000 | 3,355 |
| 1 : 4 | 19 / 76 | 171 | 471 / 1000 | 3,355 |
| independent S, C (minimum) | 20 / 57 | 157 | 471 / 892 | 3,247 |

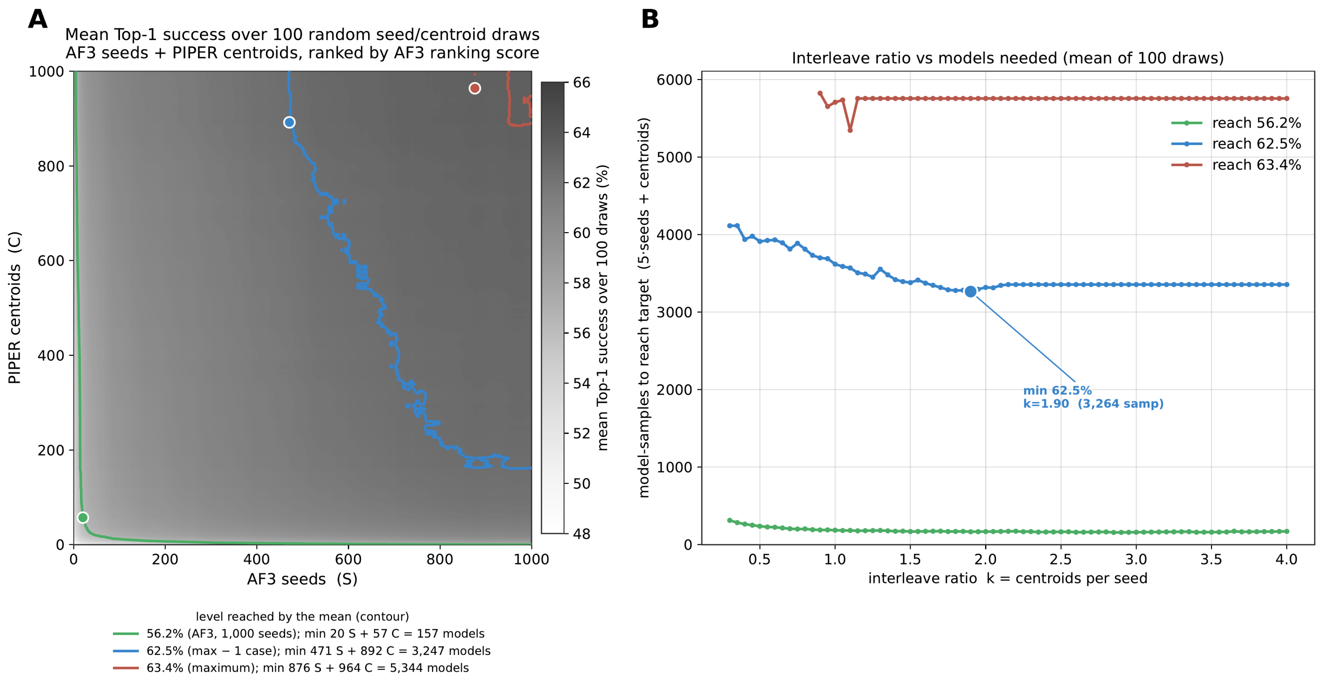


**Figure S1.** Sampling budget of the pooled method (AF3 seeds + AF3 Unmasked with PIPER templates, ranked by AF3 ranking score), averaged over 100 draws of random seed and centroid order. (A) Mean Top-1 success for every number of AF3 seeds (S) and PIPER centroids (C); contours mark where the mean reaches 56.2% (green, AF3 with 1,000 seeds), 62.5% (blue, one target below the maximum) and 63.4% (red, maximum), and dots mark the smallest budget (5S + C) reaching each level. (B) Number of models (5 per seed, 1 per centroid) at which the mean first reaches each level when seeds and centroids are added at a fixed ratio of k centroids per seed (centroids capped at 1,000). The 62.5% curve has its minimum at k = 1.9 (3,264 models). 112 targets, success defined as DockQ ≥ 0.23.

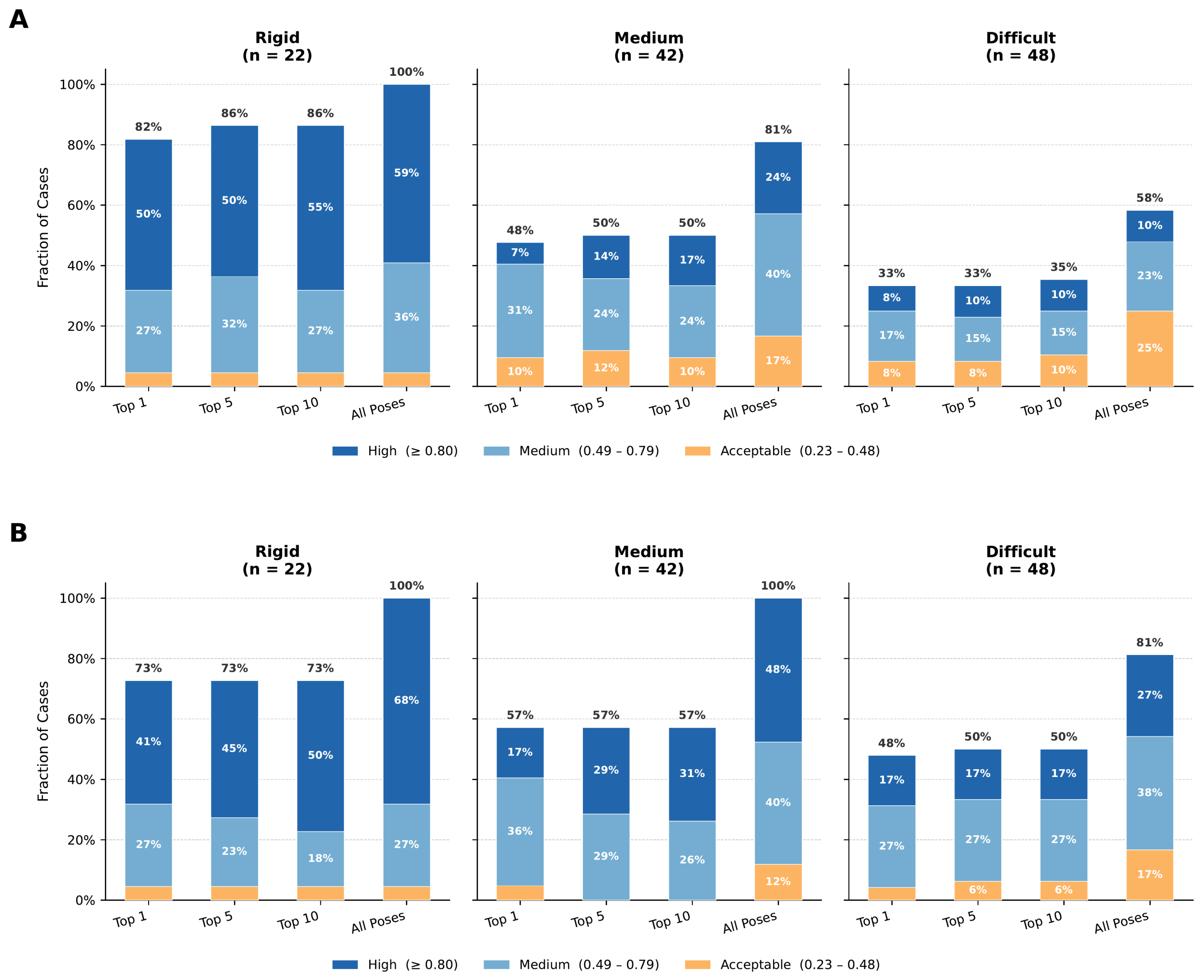


**Figure S2.** Docking templates improve antibody-antigen complex prediction by AF3 for rigid interface complexes. Success rates on the benchmark of 112 antibody–antigen complexes for baseline AF3 with unmasked PIPER templates (A) and AF3 (1,000 seeds) (B), split by interface difficulty (22 rigid, 42 medium and 48 difficult targets). Success is defined as DockQ $\geq 0.23$. Success rates for the Top-1, Top-5 and Top-10 ranked poses and for all poses (at most 1,000 PIPER centroids per target), stacked by CAPRI quality class: acceptable ($0.23 \leq$ DockQ $< 0.49$), medium ($0.49 \leq$ DockQ $< 0.80$) and high (DockQ $\geq 0.80$).

**Table S2.** Success rates of the AF3-based docking starting points, compared with baseline AF3 and the Boltz-2-based pipeline (DockQ ≥ 0.23; 112 targets; models ranked by AF3 ranking score; at most 1,000 PIPER centroids per target).

| Docking starting point | Models / target | Top-1 | Top-5 | Top-10 | All poses | Top-1 ≥ medium | Rescued / lost vs AF3 |
|---|---|---|---|---|---|---|---|
| AF3 (1,000 seeds) | 5,000 | 56.2% | 57.1% | 57.1% | 92.0% | 51.8% | — |
| Boltz-2 monomers (main text) | 1,000 | 48.2% | 50.0% | 50.9% | 75.0% | 40.2% | 11 / 20 |
| + AF3 | 6,000 | 63.4% | 67.0% | 67.0% | 93.8% | 58.0% | 10 / 2 |
| A: AF3 unbound monomers | 669 | 46.4% | 50.9% | 50.9% | 69.6% | 39.3% | 8 / 19 |
| + AF3 | 5,669 | 61.6% | 64.3% | 65.2% | 93.8% | 57.1% | 9 / 3 |
| B: split AF3 Top-1 complex | 1,000 | 58.0% | 58.0% | 58.9% | 67.0% | 53.6% | 3 / 1 |
| + AF3 | 6,000 | 58.0% | 59.8% | 60.7% | 92.0% | 53.6% | 3 / 1 |
| A + B | 1,266 | 61.6% | 62.5% | 62.5% | 78.6% | 57.1% | 8 / 2 |
| + AF3 | 6,266 | 62.5% | 64.3% | 64.3% | 93.8% | 58.0% | 8 / 1 |

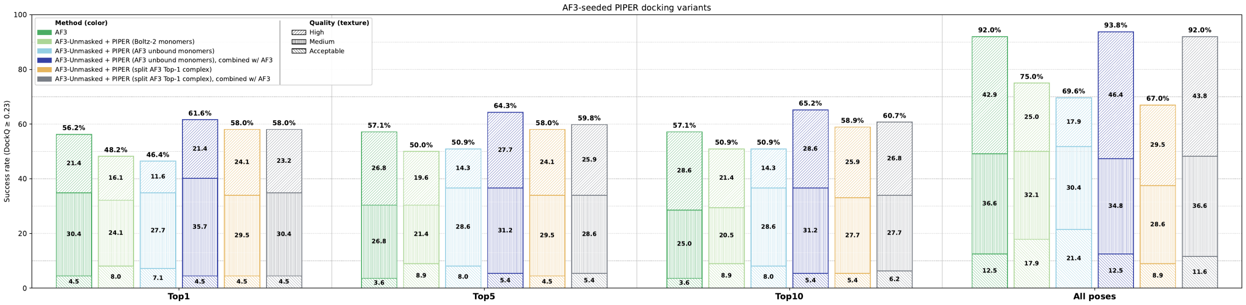


**Figure S3.** Docking AF3-generated monomers. Success rates of baseline AF3 (1,000 seeds), AF3 Unmasked with PIPER templates docked from Boltz-2 monomers (main pipeline), from AF3 unbound monomers (variant A), and from the chains of the Top-1 AF3 complex (variant B), each alone and combined with AF3. Bars show Top-1, Top-5 and Top-10 success and the full pose pool, stacked by CAPRI quality class: acceptable (0.23 ≤ DockQ < 0.49), medium (0.49 ≤ DockQ < 0.80) and high (DockQ ≥ 0.80). Models are ranked by AF3 ranking score, with at most 1,000 PIPER centroids per target; 112 targets.

**Table S3.** Score definitions and observed ranges across all four method configurations (112 targets; at most 1,000 PIPER centroids per target).

| Quantity | Boltz-2 | Boltz-2 + PIPER | AF3 | AF3-unm + PIPER |
|---|---|---|---|---|
| Score range (median) | 0.618–0.959 (0.847) | 0.707–0.968 (0.879) | 0.128–0.930 (0.569) | 0.180–0.926 (0.553) |
| sd of score | 0.056 | 0.039 | 0.172 | 0.169 |
| complex plDDT | 0.693–0.964 (0.894) | 0.773–0.972 (0.918) | — | — |
| ipTM (in formula) | 0.115–0.976 (0.664) | 0.226–0.975 (0.739) | 0.060–0.920 (0.530) | 0.100–0.890 (0.510) |
| pTM | — | — | 0.370–0.910 (0.640) | 0.400–0.910 (0.640) |
| fraction disordered | — | — | 0.000–0.270 (0.030) | 0.000–0.210 (0.020) |
| poses with has_clash | — | — | 0 | 0 |
| interface ipTM | 0.104–0.964 (0.482) | 0.118–0.966 (0.576) | 0.050–0.920 (0.170) | 0.070–0.885 (0.170) |

**Table S4.** Variance decomposition of each score into its weighted terms.

| Term | Boltz-2 | Boltz-2 + PIPER | AF3 | AF3-unm + PIPER |
|---|---|---|---|---|
| 0.8 · complex plDDT | 34.5% | 31.0% | — | — |
| 0.2 · ipTM (Boltz-2) | 40.6% | 53.1% | — | — |
| 0.8 · ipTM (AF3) | — | — | 81.5% | 78.7% |
| 0.2 · pTM | — | — | 1.7% | 1.7% |
| 0.5 · fraction disordered | — | — | 1.5% | 0.8% |
| −100 · has_clash | — | — | 0.0% | 0.0% |
| cross-terms (covariance) | 24.9% | 15.9% | 15.3% | 19.0% |

**Table S5.** Spearman ρ with DockQ — pooled (within-case).

| Quantity | Boltz-2 | Boltz-2 + PIPER | AF3 | AF3-unm + PIPER |
|---|---|---|---|---|
| complete score | 0.398 (0.066) | 0.309 (0.066) | 0.757 (0.259) | 0.686 (0.140) |
| interface ipTM | 0.407 (0.067) | 0.152 (0.059) | 0.772 (0.272) | 0.660 (0.132) |
| ρ(interface ipTM, score) | 0.672 | 0.719 | 0.717 | 0.661 |

**Table S6.** Top-1 success ranked by model score versus by interface ipTM, against the oracle.

| **Top-1 success (DockQ ≥ 0.23)** | Boltz-2 | Boltz-2 + PIPER | AF3 | AF3-unm + PIPER |
|---|---|---|---|---|
| ranked by model score | 12.5% | 17.9% | 56.2% | 48.2% |
| ranked by interface ipTM | 16.1% | 11.6% | 55.4% | 45.5% |
| difference | +3.6 pts | −6.2 pts | −0.9 pts | −2.7 pts |
| oracle | 62.5% | 76.8% | 92.0% | 75.0% |
| **Top-1 success (DockQ ≥ 0.49)** | | | | |
| ranked by model score | 5.4% | 10.7% | 51.8% | 40.2% |
| ranked by interface ipTM | 7.1% | 7.1% | 50.9% | 41.1% |
| oracle | 37.5% | 48.2% | 79.5% | 57.1% |

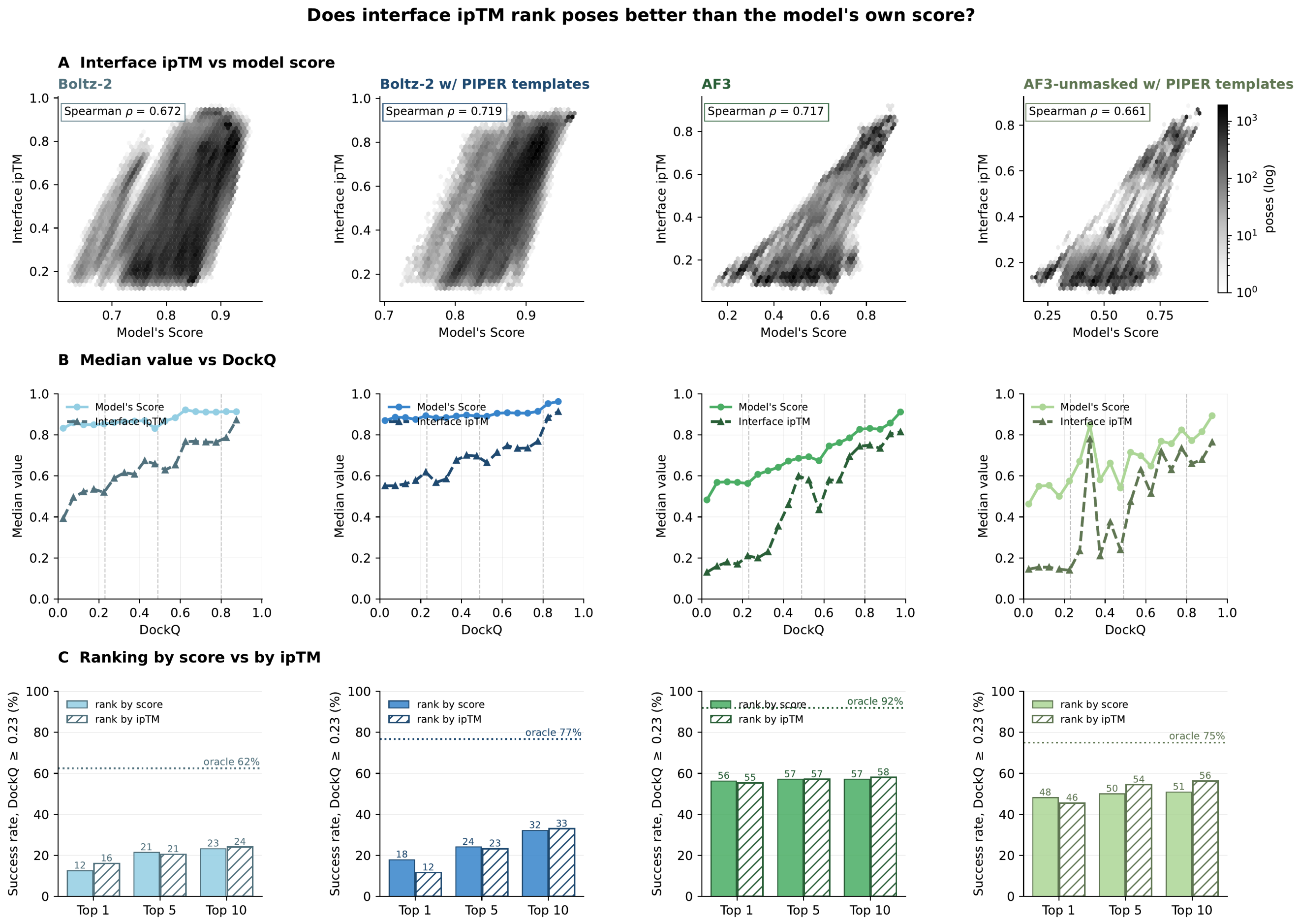


**Figure S4.** Interface ipTM versus the model's own score. (A) Joint density per method with Spearman ρ. (B) Median value versus DockQ for both quantities. (C) Top-N success ranking by score versus by ipTM, against the oracle.

**Table S7.** Success rates of Boltz-2 with the per-chain templates retrieved by AF3 (DockQ ≥ 0.23; 111 targets; models ranked by Boltz-2 confidence score).

| Method | Models / target | Top-1 | Top-5 | Top-10 | All poses | Top-1 ≥ medium | Rescued / lost vs Boltz-2 (100 seeds) |
|---|---|---|---|---|---|---|---|
| Boltz-2 (100 seeds) | 500 | 13.5% | 21.6% | 24.3% | 45.9% | 5.4% | — |
| Boltz-2 + AF3 templates (same 100 seeds) | 500 | 18.0% | 24.3% | 25.2% | 49.5% | 9.0% | 7 / 2 |
| pooled | 1,000 | 16.2% | 24.3% | 26.1% | 54.1% | 8.1% | 4 / 1 |
| Boltz-2 (1,000 seeds) | 5,000 | 11.7% | 20.7% | 22.5% | 62.2% | 5.4% | 1 / 3 |

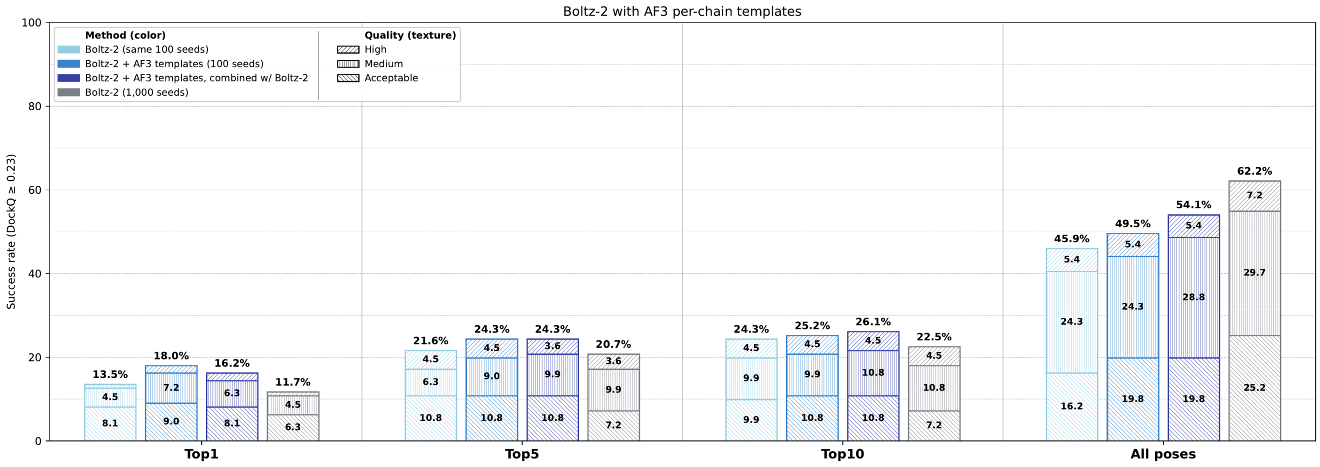


**Figure S5.** Boltz-2 with the per-chain templates retrieved by AF3. Success rates of untemplated Boltz-2 and Boltz-2 with AF3 templates on the same 100 seeds, the two pooled, and untemplated Boltz-2 with 1,000 seeds, shown for the Top-1, Top-5 and Top-10 ranked models and the full pose pool and stacked by CAPRI quality class as in Fig. S3; 111 targets.

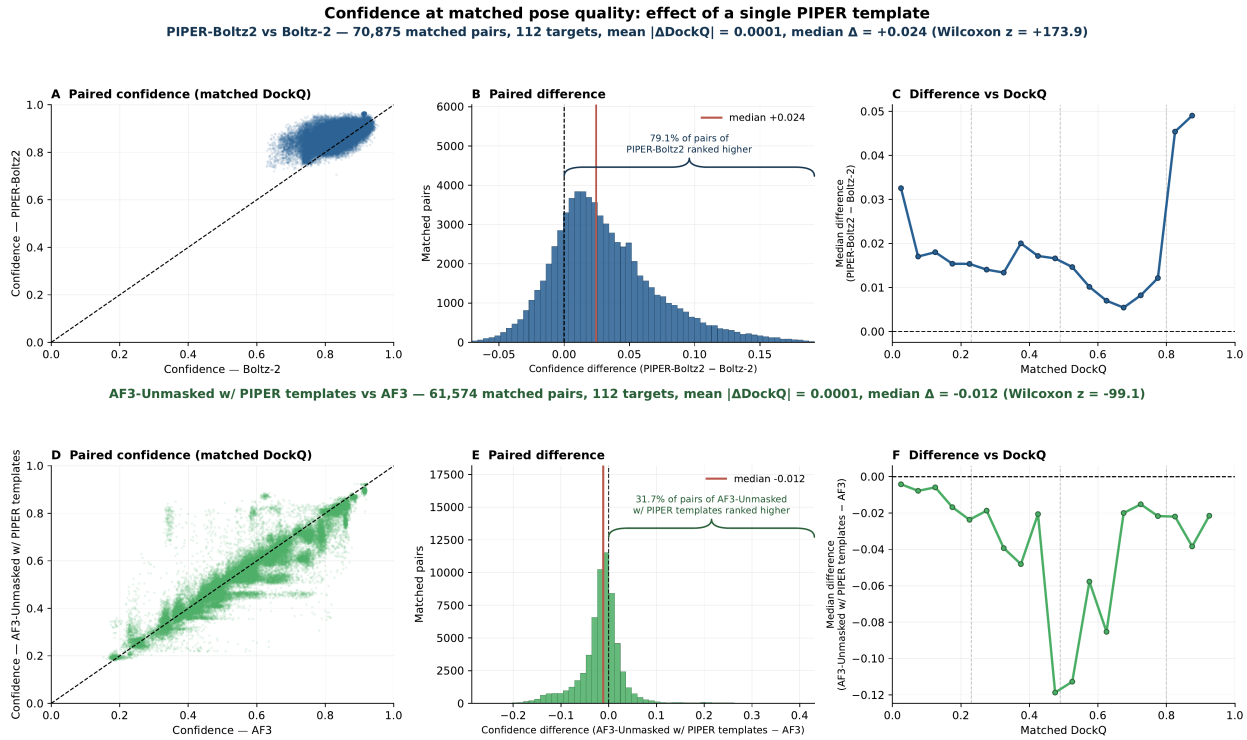


**Figure S6.** Confidence at matched pose quality. Poses were paired within each target by DockQ (≤ 0.005 apart). (A–C) PIPER-Boltz2 vs Boltz-2, 70,875 pairs; (D–F) AF3-Unmasked w/ PIPER templates vs AF3, 61,574 pairs; 112 targets each, at most 1,000 PIPER centroids per target. Panels show paired confidences with the identity line (A, D), the distribution of differences with the median marked and a brace giving the fraction of pairs in which the templated run scored higher (B, E), and the median difference across DockQ bins (C, F).

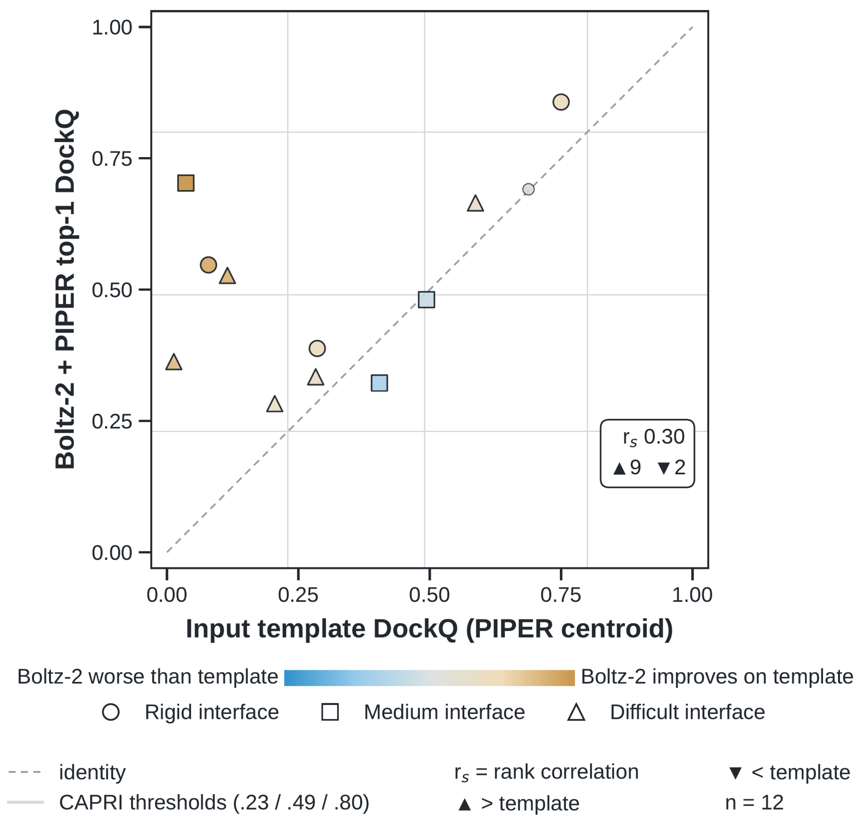


**Figure S7.** DockQ of the Top-1 Boltz-2 + PIPER model versus that of the PIPER centroid used as its template, for the 12 targets in which untemplated Boltz-2 fails at Top-1 (DockQ < 0.23) and Boltz-2 with PIPER templates succeeds. Colour gives the change in DockQ (gold, Boltz-2 model higher than the template; blue, lower) and marker shape the interface difficulty class. Dashed line, identity; grey lines, CAPRI thresholds (0.23, 0.49, 0.80). rs, Spearman rank correlation; ▲ and ▼, number of targets above and below the template ($|\Delta\text{DockQ}| \leq 0.005$ counted as a tie); 112-target benchmark.
